# Metabolic Trajectories During Surgical Stress in Patients Undergoing Cardiac Surgery

**DOI:** 10.1101/2025.09.16.676529

**Authors:** Dohyun Ku, Jackson Bartelt, Jenna Feeley, M. Citlalli Perez-Guzman, Lizda Guerrero-Arroyo, Amalia Abraham, Saaki Kollipara, Nicolas Gonzalez, Sabeena Usman, Helaina E. Huneault, Andrea Corujo-Rodriguez, Georgia M. Davis, Richard G. Kibbey, Dean P. Jones, Thomas R. Ziegler, Michael Halkos, Arshed A. Quyyumi, M. Ryan Smith, Jing Li, Francisco J. Pasquel

## Abstract

Stress hyperglycemia (SH) during acute illness is linked to adverse surgical outcomes, yet the accompanying metabolic perturbations are incompletely characterized. We profiled longitudinal metabolic changes in adults without diabetes undergoing cardiac surgery to identify pathways associated with perioperative SH (defined as point-of-care glucose ≥140 mg/dL on ≥3 readings or ≥180 mg/dL once). Blood was collected at baseline before surgery (T0) and at 2 h (T1), 24–48 h (T2), and 72–96 h (T3) after surgical initiation. High-resolution metabolomics (LC–MS) was integrated with continuous glucose monitoring and inflammatory/cardiac biomarkers. At T0, several pathways were associated with subsequent SH, including bile acid metabolism, the carnitine shuttle, and fatty-acid oxidation, suggesting preoperative metabolic susceptibility. In longitudinal analyses, participants who developed SH showed coordinated postoperative changes with significant enrichment of pathways not evident at baseline, C21-steroid hormone biosynthesis, glycerophospholipid metabolism, and glycosphingolipid (ceramide) metabolism, consistent with lipid remodeling and inflammatory signaling during surgical stress. Individuals with SH also exhibited higher inflammatory biomarker levels (high-sensitivity C-reactive protein and soluble urokinase plasminogen activator receptor). A machine-learning model using early metabolomic features predicted SH with an area under the receiver operating characteristic curve of 0.86. These findings highlight distinct preoperative and perioperative metabolic trajectories associated with SH and implicate established dysglycemia-related pathways, as well as stress-induced pathways in perioperative metabolic dysregulation. Pathway enrichment analyses were exploratory and hypothesis-generating; validation in larger cohorts and assessment of implications for clinical outcomes are warranted.

## INTRODUCTION

Glucose is the most routinely assayed metabolite in clinical medicine, and it’s widely used to gauge metabolic stability in patients with and without diabetes. However, during acute physiological stress such as major surgery, hyperglycemia may signal a much deeper biochemical disruption. Perioperative glucose elevations are a recognized hallmark of surgical stress and correlate with higher rates of surgical-site infection, impaired wound healing, prolonged hospital stay, and death.^1^ However, the broader metabolic derangements that accompany glucose excursions, as well as the mechanistic links between hyperglycemia and poor outcomes, remain inadequately defined.

Coronary-artery bypass grafting (CABG) is a well-standardized, high-intensity surgical procedure that evokes reproducible metabolic and inflammatory responses. Stress hyperglycemia (SH) occurs commonly in CABG patients and is consistently associated with adverse events,^2^ yet the metabolic pathways driving this response, and their clinical relevance remain largely unknown. Since CABG follows a predictable operative timeline, it offers a unique in-vivo model for studying the temporal evolution of stress-induced metabolic changes.

Recent advances in high-resolution metabolomics, continuous glucose monitoring (CGM), and integrative bioinformatics enable simultaneous tracking of thousands of metabolites alongside real-time glycemic profiles. Here, we applied longitudinal plasma metabolomics to detect changes in individual metabolites, CGM to characterize glucose changes, and targeted biomarker assays to determine if metabolites and biomarkers fluctuated with surgical stress and SH among adults without diabetes undergoing CABG. Using pathway enrichment analysis and machine learning classification, we defined metabolic trajectories linked to perioperative SH and delineated the biochemical networks that couple stress-induced glucose rises to inflammatory and metabolic pathways.

## RESULTS

Figure 1 provides an overview of the study design and analytical workflow. Further methodological details are provided in the Methods section.

**Figure 1.**
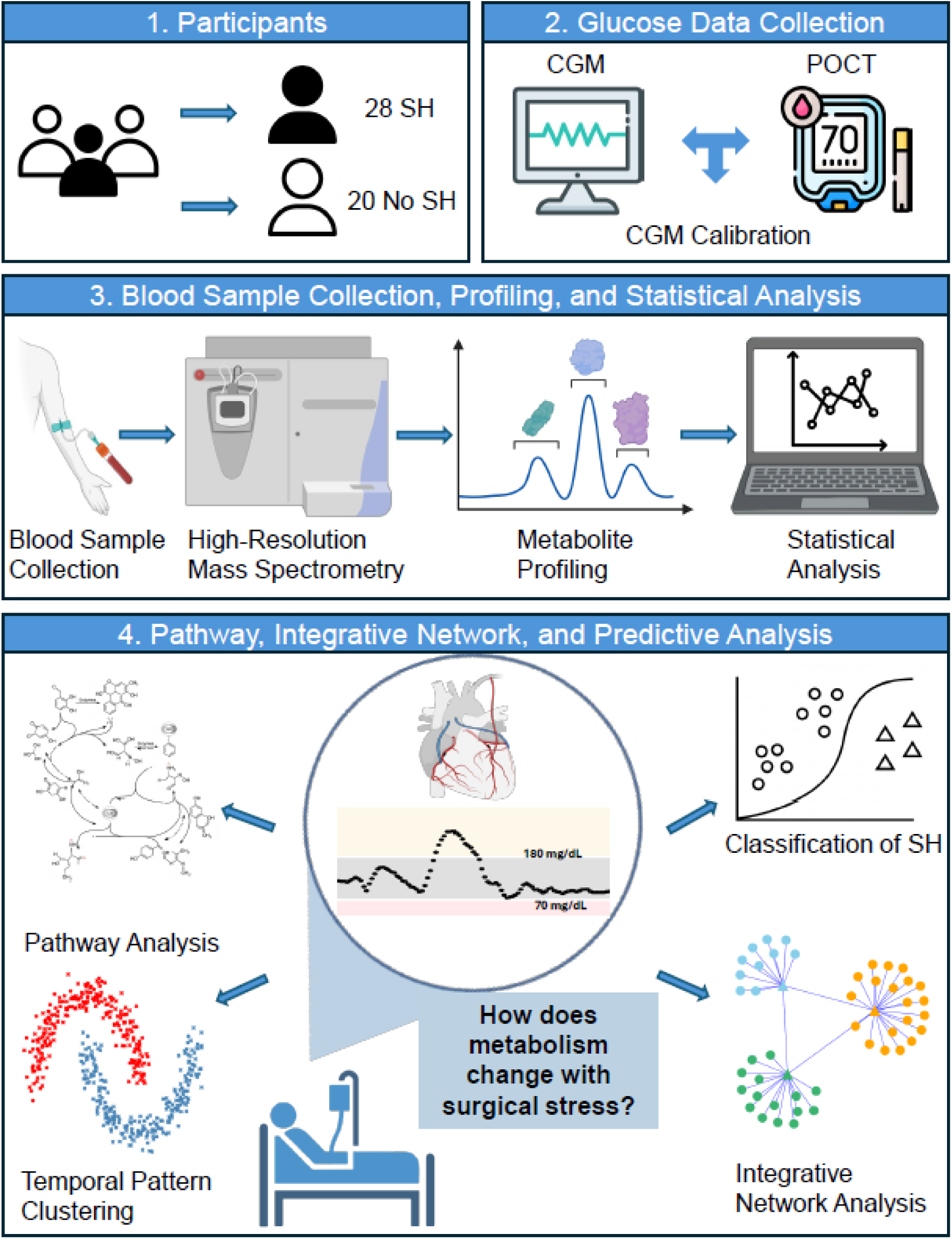
Flow of study methods. A total of 48 participants undergoing CABG surgery were enrolled, including 28 with and 20 without stress hyperglycemia (SH). 2. Glucose levels were monitored throughout the perioperative period using point-of-care testing (POCT) and continuous glucose monitoring (CGM) devices. CGM readings were calibrated using POCT measurements. 3. Blood samples were collected at baseline, 2, 24–48, and 72–96 hours after surgery initiation. These samples were analyzed using high-resolution mass spectrometry to profile metabolites and biomarkers. Nonlinear mixed-effects models were applied to identify metabolites with significant temporal changes across the perioperative period. 4. Significant metabolites were subjected to pathway enrichment analysis, and their temporal trajectories were clustered to reveal dominant dynamic patterns. A participant-level classifier was developed using selected metabolites to predict SH. Integrated network analysis including community detection was performed to identify variable clusters associated with SH. Key metabolic pathways and metabolites were further evaluated for biological relevance to surgical stress and hyperglycemia.

### Participant Characteristics

A total of 48 participants with metabolomics data were included in the analysis [mean age = 64.0 ± 10.1 years; 42% male; body mass index (BMI) = 28.3 ± 4.5 *Kg*/*m*^2^]. Among these participants, 28 (58%) developed SH. **Supplementary Table 1** presents the demographic characteristics of study participants.

### Metabolites at Baseline (before Surgery) Associated with Stress Hyperglycemia

At baseline, two-sample t-tests were performed to compare SH and non-SH participants, identifying 602 and 769 significant putative metabolites using reverse-phase chromatography (C18) and Hydrophilic Interaction Liquid Chromatography (HILIC), respectively. Subsequent pathway enrichment analysis identified 22 significant metabolic pathways via C18 and 17 via HILIC (see **Supplementary Table 2**).

We applied a machine learning pipeline to predict SH using baseline metabolite levels identified from pathway enrichment analysis. This pipeline with logistic regression demonstrated the best predictive performance (AUC of 0.79, accuracy of 0.75, sensitivity of 0.79, and specificity of 0.70). ROC curves for all four methods are shown in Figure 2A. Comparative performance results for all models are presented in **Supplementary Table 3**. Figure 2B presents the top 20 metabolites ranked by feature importance in the pipeline. Detailed information on m/z values, retention times, annotations, and abbreviations is provided in **Supplementary Table 4**.

**Figure 2A-C.**
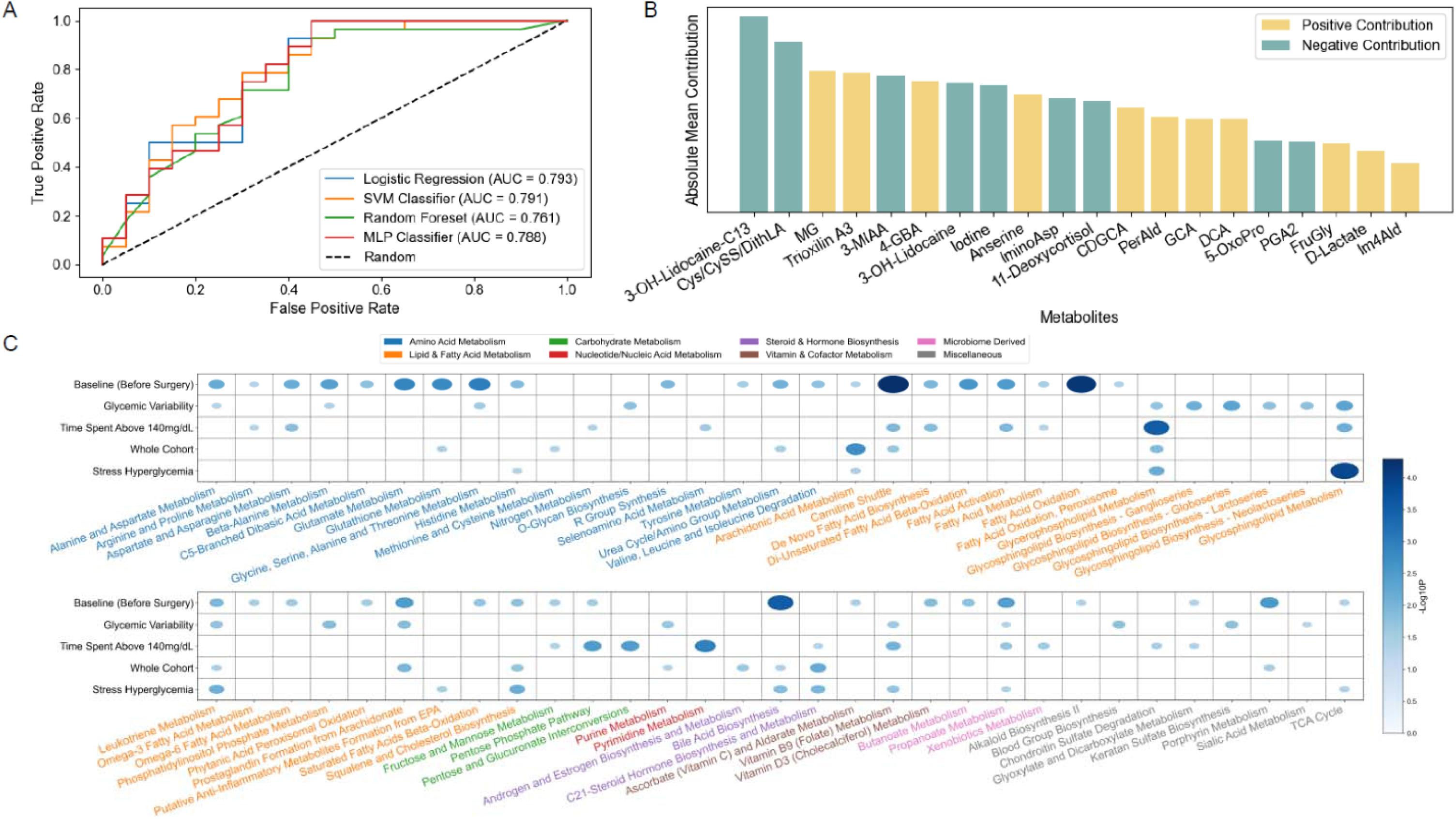
SH prediction by baseline metabolites and pathway enrichment analysis across perioperative period. **(A)** Area under the receiver operating characteristic curves (AUCs) comparing four machine learning models for stress hyperglycemia (SH) prediction using baseline metabolites. Machine learning pipeline with logistic regression achieved the highest performance (AUC = 0.793). **(B)** Feature importance ranking of the top 20 metabolites using baselin metabolites. Bar heights represent absolute mean contribution to model predictions, with metabolites color-coded based on their contribution type (positive contribution in orange, negative contribution in teal). **(C)** Pathway enrichment analysis showing the most strongly associated metabolic pathways across five different analyses: 1) metabolites at baseline associated with SH, 2) metabolites associated with time above range (>140 mg/dL), 3) metabolites associated with glycemic variability (coefficient of variation), 4) metabolites associated with temporal changes for whole cohort, and 5) metabolites associated with SH across four time points. Bubble size and color intensity indicate correlation strength (-log10 p-value, scale 0-5).

### Significant Pathways Associated with Time Above Range (>140 mg/dL) and Glycemic Variability (Coefficient of Variation)

We investigated temporal metabolomic patterns in the whole cohort using all four time points: 24 hours before surgery (T0), 2 hours after surgery initiation (T1), 24-48 hours after surgery initiation (T2), and 72-96 hours after surgery initiation (T3). To identify metabolic pathways associated with glycemic variability and hyperglycemic exposure during the 48 hours following surgery initiation, we performed clinical difference analyses using two continuous variables: coefficient of variation (CV) and time above range (TAR, >140 mg/dL). To obtain complete CGM traces for the 48-hour after surgery, we imputed missing readings and calibrated CGM values with concurrent POCT results. Calibration results and examples are provided in the Supplementary Section. From the resulting data, we calculated CV and TAR, each used as a continuous interacting predictor with time in the nonlinear mixed-effects model (NMEM) for every metabolite across four time points. Figure 3A reveals a clear separation between groups, with participants experiencing SH consistently showing higher glucose levels in the perioperative period. This difference gradually diminishes until approximately 40 hours after surgery initiation.

**Figure 3 A-D.**
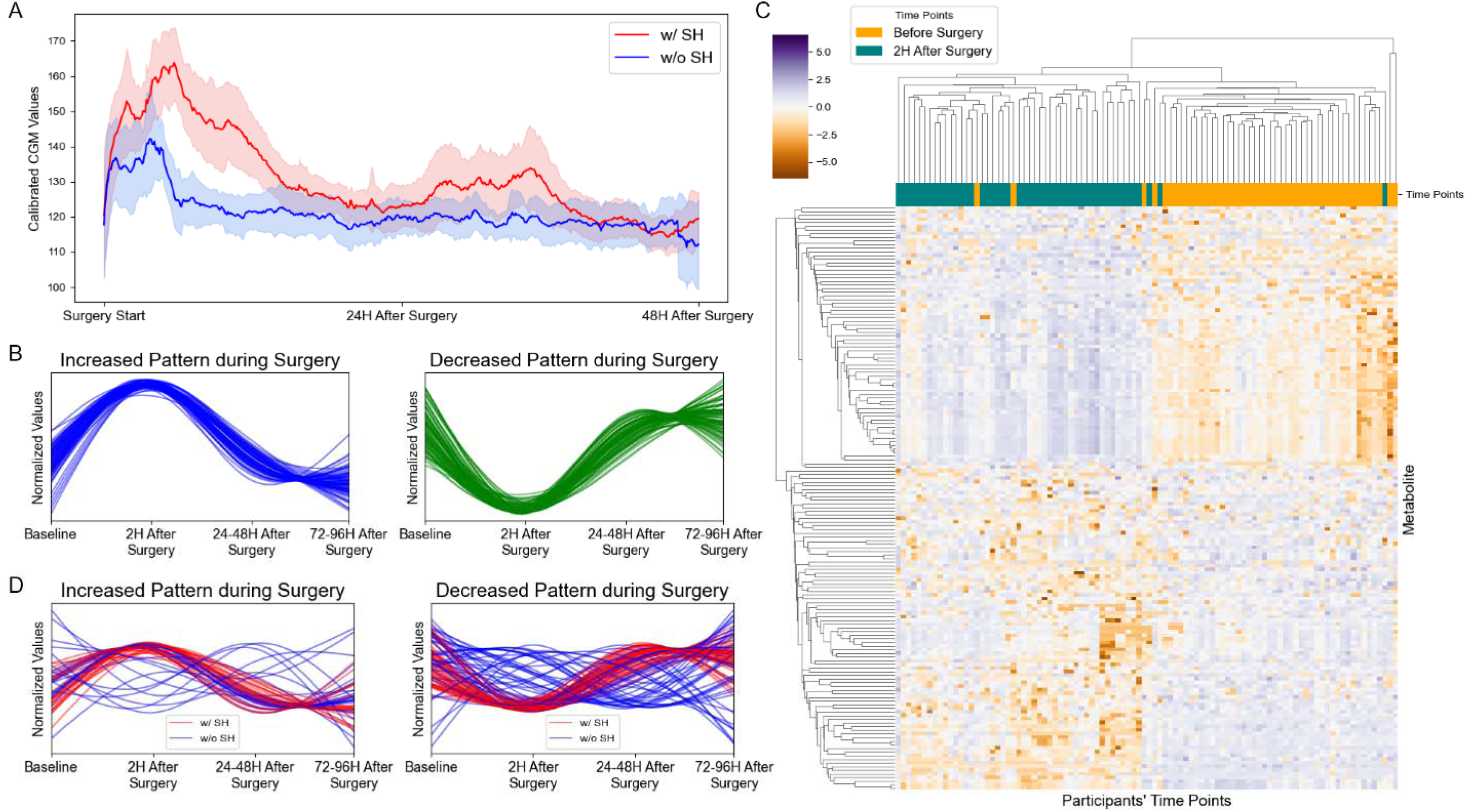
Glucose profiles and metabolic trajectories during perioperative period. **(A)** shows average CGM profiles comparing participants with stress hyperglycemia (SH) (red line) versus those without (blue line) from surgery start through 48 hours after surgery start. **(B)** shows the dominant metabolite temporal patterns identified from nonlinear mixed-effects modeling in the whole cohort. The left panel displays metabolites with increased levels during surgery with subsequent return to baseline; right panel shows metabolites with decreased levels during surgery with later recovery. All metabolite values are normalized. **(C)** Heatmap visualization of metabolite changes between baseline and 2 hours after surgery start. Hierarchical clustering reveals distinct metabolite groups based on temporal patterns. Color scale indicates normalized metabolite levels from lower levels (purple) to higher levels (orange). **(D)** Group-specific metabolite trajectories comparing participants with SH (red lines) versus those without (blue lines). Left panel shows metabolites with surgery-induced increases; right panel indicates metabolites with surgery-induced decreases. Participants without hyperglycemia did not show a dominant temporal pattern. Metabolite levels are normalized.

After applying subsequent pathway enrichment analysis, we identified several putative metabolites associated with CV (1,628 in C18 and 1,667 in HILIC) and TAR (991 in C18 and 1,115 in HILIC). Pathways with the highest association with glycemic variability by CV included glycosphingolipid metabolism, glycerophospholipid metabolism, purine metabolism, vitamin B9 (folate) metabolism, glycine, serine, alanine and threonine metabolism, glycosphingolipid metabolism, and phosphatidylinositol phosphate metabolism. **Supplementary Table 5** shows all significant pathways associated with CV along with overlap and pathway size. Pathways with the highest association with TAR included pyrimidine metabolism, pentose phosphate pathway, pentose and glucuronate interconversions, vitamin B9 (folate) metabolism, aspartate and asparagine metabolism, glycerophospholipid metabolism, glycosphingolipid metabolism, carnitine shuttle, and fatty acid activation. **Supplementary Table 6** shows all significant pathways associated with TAR including overlap and pathway size. Figure 2C shows pathway enrichment analyses showing the most strongly associated pathways with glycemic variability (CV) and time spent above 140 mg/dL (TAR).

### Dominant Temporal Metabolic Trajectories during Surgical Stress

Based on all four time points in the whole cohort, NMEM identified 5,123 and 6,500 significant mass spectrometry features showing significant temporal variation using C18 and HILIC, respectively. Several metabolic pathways (e.g. arachidonic acid metabolism, C21-steroid hormone biosynthesis and metabolism, glycerophospholipid metabolism, carnitine shuttle, bile acid metabolism) were identified via C18 and HILIC (**Supplementary Table 7** shows all significant pathways including significance and overlap/pathway size). Figure 2C shows pathway enrichment analyses showing the most strongly associated pathways with surgical stress in the whole cohort.

All putative metabolites from significant pathways (n=313) were combined, and metabolite trajectories were extracted from the NMEM full model in the whole cohort statistical analysis, identifying a single, predominant *“early-spike/rapid-recovery”* trajectory that accounted for 35.8% of all profiled compounds (86 detected on the C18 column and 74 on the HILIC column). Metabolite intensities in this cluster rose sharply from baseline, peaking 2 hours after surgery, followed by rapid decline to baseline or slightly below within 24–48 hours, with levels remaining stable through 72–96 hours, shown in Figure 3B. Consistent with the dominant clustering pattern, the Figure 3C heatmap highlights metabolites that undergo significant up- and down-regulation at two hours after surgery initiation.

### Dominant Metabolomic Temporal Patterns from Significant Pathways Associated with Hyperglycemia

To determine whether metabolomic temporal patterns varied by glycemic status, we applied NMEM using SH as a binary group variable. This analysis identified 1,733 significant putative compounds via C18 and 1,353 via HILIC. Subsequent pathway enrichment analysis revealed several significant metabolic pathways in both C18 and HILIC, as shown in **Supplementary Table 8 and** Figure 2C.

Multiple putative compounds were detected, resulting in the identification of 78 metabolites via C18 and 39 via HILIC. A density-based spatial clustering of applications with noise (DBSCAN) algorithm identified a dominant temporal pattern in participants with SH that encompassed 62.4% of the identified metabolites, containing 48 metabolites from C18 and 25 metabolites from HILIC. The pattern also showed significant metabolite level changes at T1 (2 hours) with a subsequent return to the original level at T2 (72-96 hours). In contrast, participants without SH did not exhibit a clear dominant pattern, as illustrated in Figure 3D.

### Metabolomic Changes during Surgery Differentiate Groups with and without Stress Hyperglycemia

To identify metabolites that can distinguish SH versus non-SH participants, both the whole cohort and group difference analyses were subjected to further comparative analysis. Two sample t-tests were performed to compare the changes in metabolite levels between T0 and T1, representing the transition from the baseline to early intraoperative phase (*during CABG*) in participants with and without SH.

From the whole cohort analysis (*without* SH as a binary group variable) out of 160 putative metabolites, 22 achieved statistical significance (Figure 4A), with all metabolites showing greater absolute changes (baseline to 2 hours) among participants with SH. Similarly, group difference analysis of 73 metabolites identified 17 statistically significant metabolites (Figure 4B), of which 15 exhibited greater absolute changes in the SH group. Complete metabolite details, including m/z values, retention times, annotations, abbreviation, and clinical relevance, are provided in **Tables 1** and **2**, respectively. These findings indicate that specific metabolite changes between T0 and T1 may serve as predictive biomarkers for SH development.

**Figure 4 A-D.**
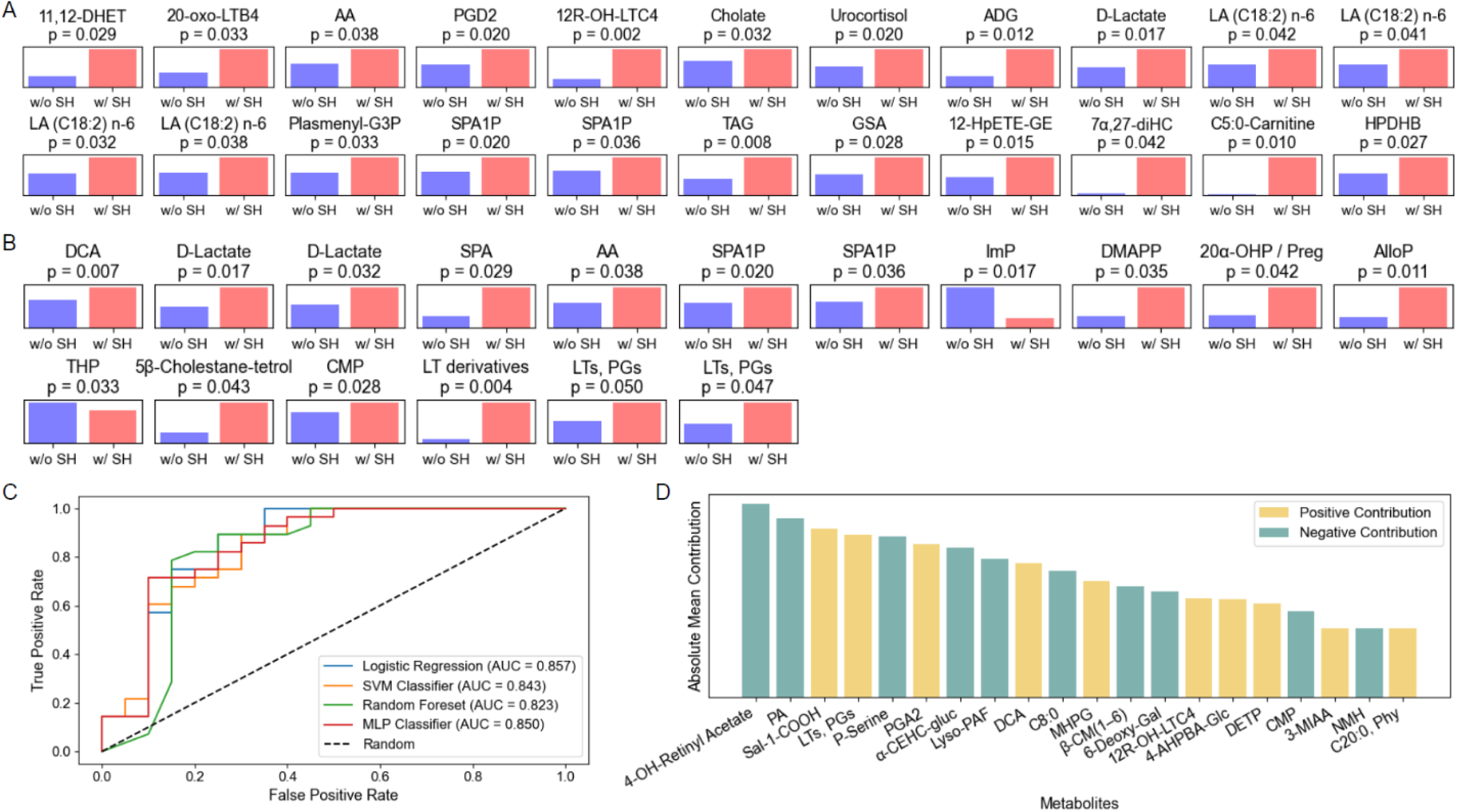
Metabolic profiles for SH versus non-SH participants and SH prediction. (**A)** Bar plots showing 22 metabolites from whole cohort analysis with statistically significant differences in temporal changes (T0 to T1) between participants with stress hyperglycemia (SH) (red bars) and without (blue bars). Each subplot displays mean absolute metabolite changes with p-values from two-sample t-tests. All 22 metabolites showed greater absolute changes in the SH group. **(B)** Bar plots showing 17 metabolites from group difference analysis with significant group differences in temporal changes. Among these, 15 metabolites (all except ImP and THP) exhibited greater absolute changes in the participants with **SH**. **(C)** Area under the receiver operating characteristic curves (AUCs) comparing four machine learning models for **SH** prediction. Machine learning pipeline with logistic regression achieved the highest performance (AUC = 0.857). **(D)** Feature importance ranking of the top 20 metabolites. Bar heights represent absolute mean contribution to model predictions, with metabolites color-coded based on their contribution type (positive contribution in orange, negative contribution in teal).

**Table 1.** Significantly Altered Metabolites from Whole Cohort Analysis.

| Column | Mummichog pathway Label | m/z_RT | Adducts | Mummichog, KEGG Label | Abbreviation | Patterns | Relevance |
| --- | --- | --- | --- | --- | --- | --- | --- |
| C18 | Arachidonic acid metabolism | 337.2359_243.3 | $[M-2H]^{2-}$ | 11,12-DHET | - | Increased | An inverse relationship has been reported between plasma 11,12-DHET and glucose.<sup>41</sup> Mice with diabetes exhibit higher levels of 11,12-DHET.<sup>42</sup> |
| | | 348.1928_205.8 | $[M-H_2O-H]^-$ | 20-oxo-leukotriene B4 | 20-oxo-LTB4 | Increased | Leukotriene B4 is produced in higher levels in those with diabetes and has been associated with poor wound healing.<sup>43,4</sup> |
| | | 349.2381_225.6 | $[M-H]^-$ | Arachidonic acid | AA | Increased | Increased AA levels decrease or inhibit insulin release.<sup>6</sup> In addition, PGF2 is elevated in mice with diabetes.<sup>5</sup> |
| | | 351.2207_242.5 | $[M+^{13}C-H]^-$ | Prostaglandin D2 | PGD2 | Increased | PGD2 can improve insulin resistance,<sup>44</sup> and reduce inflammation during a myocardial infarction.<sup>3</sup> |
| | | 677.2446_148.7 | $[M-H]^-$ | 10,11-dihydro-12R-hydroxy-leukotriene C4 | 12R-OH-LTC4 | Increased | Leptin level was found to be inversely correlated with levels of 10,11-dihydro-12R-hydroxy-leukotriene.<sup>45</sup> In addition, elevated Leukotriene A4 Hydrolase in the Leukotriene pathway were found to be associated with cardiac death.<sup>46</sup> |
| | Bile acid biosynthesis | 203.1359_24.6 | $[M(C^{13})-H]^-$ | Cholate | - | Decreased | Cholate can prevent a hyperglycemic response to a high fat diet.<sup>47</sup> Bile acids have been used to improve glucose regulation.<sup>48</sup> |
| | C21-steroid hormone biosynthesis and metabolism | 347.2204_228.4 | $[M-2H]^{2-}$ | Urocortisol | - | Increased | Urocortisol is a breakdown product of cortisol,<sup>49</sup> a potent anti-inflammatory hormone. Cortisol is elevated after cardiac surgery.<sup>49</sup> Higher cortisol is associated with lower percent time in target glucose range.<sup>50</sup> |
| | | 466.2482_274.8 | $[M-H]^-$ | Androsterone glucuronide | ADG | Decreased | When adjusted for age, education, race/ethnicity, smoking, and alcohol use, androstanediol glucuronide is associated with worse cardiovascular risk factors and death due to heart disease.<sup>51</sup> |
| | Glycosphingolipid metabolism | 380.2556_268.8 | $[M-2H]^{2-}$ | Sphinganine 1-phosphate | SPA1P | Increased | Sphinganine-1-P is converted to sphingosine-1-P which lowers blood glucose levels and is therefore protective after cardiac surgery.<sup>7</sup> Increased Sphinganine-1-P may represent lack of conversion to Sphingosine-1-phosphate. |
| | | 381.2569_268.2 | $[M-H]^-$ | | | Increased | |
| | Glycerophospholipid metabolism | 130.0509_19.2 | $[M-H]^-$ | D-Lactate | - | Decreased | Lactate levels are intimately tied to glycolytic flux and hyperglycemia.<sup>52</sup> Lactate levels also increase after surgery and in those with hyper or hypoglycemia.<sup>52</sup> Cardiac failure is also associated with D-lactic acidosis.<sup>53</sup> |
| | | 279.2328_229.7 | $[M+^{13}C-H]^-$ | linoleic acid (all cis C18:2) n-6 | LA (C18:2) n-6 | Increased | LA is beneficial for cardiovascular health; that is, higher LA intake is associated with lower CVD risk.<sup>54</sup> Better glucose management was also observed with LA supplementation.<sup>55</sup> |
| | | 280.2361_229.2 | $[M-H]^-$ | | | Increased | |
| | | 325.2383_229.8 | $[M-H]^-$ | | | Increased | |
| | | 339.2538_229.2 | $[M+^{13}C-H]^-$ | | | Increased | |
| | | 327.2327_219.3 | $[M-H+O]^-$ | 2-Acyl-1-(1-alkenyl)-sn-glycero-3-phosphate | Plasmenyl-G3P | Increased | Lysophosphatidic acid (LPA) (a glycerophospholipid) impairs insulin release and therefore impairs glucose tolerance.<sup>56</sup> After a myocardial infarction, LPA is also involved in cardiac dysfunction and hypertrophy.<sup>57</sup> |
| | | 380.2556_268.8 | $[M-2H]^{2-}$ | Sphinganine 1-phosphate | SPA1P | Increased | Sphinganine-1-P is converted to sphingosine-1-P which lowers blood glucose levels and is therefore protective after cardiac surgery.<sup>7</sup> Increased Sphinganine-1-P may represent lack of conversion to Sphingosine-1-phosphate. |
| | | 381.2569_268.2 | $[M-H]^-$ | | | Increased | |
| | | 637.539_254.7 | $[M-H_2O-H]^-$ | Triacylglycerol | TAG | Increased | Elevated triacylglycerols are seen in those with myocardial infarction.<sup>58</sup> High triglycerides increase insulin resistance, and breakdown products (glycerol + free fatty acids) stimulate gluconeogenesis.<sup>59</sup> |
| | Porphyrim metabolism | 171.0775_17.0 | $[M+HCOO]^-$ | L-Glutamate 5-semialdehyde | GSA | Decreased | Overall, glutamate, but not specifically L-Glutamate 5-semialdehyde, accumulated in higher levels in the brain for patients with diabetes. <sup>60</sup> |
| | Prostaglandin formation from arachidonate | 391.2519_259.2 | $[M-H]^-$ | 12-hydroperoxyeicosa tetraenoate glyceryl ester | 12-HpETE-GE | Increased | Prostaglandins help regulate glucose metabolism and keep blood glucose levels in a healthy range.<sup>61</sup> Prostaglandins are also implicated in atherosclerosis and myocardial infarction.<sup>62</sup> |
| HILIC | Bile acid biosynthesis | 417.3368_25.5 | $[M-H]^-$ | 7-α,27-Dihydroxycholesterol | 7α,27-diHC | Decreased | Oxysterols, such as 7βHC and 7keto-cholesterol, have cytotoxic effects and have been found to be linked with inflammation and general metabolic imbalance.<sup>63</sup> |
| | Carnitine shuttle | 384.3106_24.4 | $[M-H_2O-H]^-$ | Pentadenoyl carnitine | C5:0-Carnitine | Decreased | Supplementation with L-carnitine reduces glucose levels.<sup>64</sup> Supplementation can help with glucose regulation when insulin resistance is present.<sup>64</sup> |
| | Urea cycle/amino group metabolism | 543.3793_25.2 | $[M-H_2O-H]^-$ | 3-Hexaprenyl-4,5-dihydroxybenzoate | HPDHB | Increased | Arginine is elevated and ornithine is decreased in those with T2D.<sup>65</sup> Lower insulin sensitivity and glucose tolerance have been correlated with urea.<sup>65</sup> |
| Twenty-two metabolites identified from whole cohort analysis showing statistically significant temporal changes between pre-surgery baseline (T0) and 2 hours after surgery initiation (T1) across all participants. Metabolites are organized by metabolic pathways identified through Mummichog. For each metabolite, the table provides chromatographic details (column type, m/z, retention time), adducts, annotations from Mummichog and KEGG, abbreviations, direction of change (increased/decreased), and clinical relevance with supporting references. |  |  |  |  |  |  |  |

**Table 2.** Significantly Altered Metabolites from Stress Hyperglycemia Group Difference Analysis.

| Column | Mummichog Pathway Label | m/z_RT (s) | Adducts | Mummichog, KEGG Label | Abbreviation | Patterns | Relevance |
| --- | --- | --- | --- | --- | --- | --- | --- |
| C18 | Bile acid biosynthesis | 195.139_144.7 | $[M-H_2O-H]^-$ | Deoxycholic acid | DCA | Increased | Bile acid metabolism leads to glucose perturbation via lipid dysregulation & direct stimulation of nuclear & membrane bound receptors.<sup>66</sup> |
| | Glycerophospholipid metabolism | 130.0509_19.2 | $[M-H]^-$ | D-Lactate | - | Decreased | Lactate levels are intimately tied to glycolytic flux and hyperglycemia.<sup>52</sup> Lactate levels also increase after surgery and in those with hyper or hypoglycemia.<sup>52</sup> Cardiac failure also shown to be associated with D-lactic acidosis.<sup>53</sup> |
| | | 149.0455_21.9 | $[M-H]^-$ | | | Decreased | |
| | | 337.2359_243.3 | $[M-H]^-$ | Sphinganine | SPA | Increased | Sphinganine may be related to hyperglycemia and insulin resistance.<sup>8</sup> |
| | | 349.2381_225.6 | $[M-H]^-$ | arachidonic acid | AA | Increased | See Table 1 |
| | | 380.2556_268.8 | $[M-H_2O-H]^-$ | Sphinganine-1-phosphate | SPA1P | Increased | See Table 1 |
| | | 381.2569_268.2 | $[M-H]^-$ | | | Increased | |
| | Histidine metabolism | 155.0462_90.0 | $[M-H_2O-H]^-$ | 4-Imidazolone-5-propanoate | ImP | Decreased | Imidazole propanoate has been associated with insulin resistance, T2D, and heart failure.<sup>67</sup> Histidine itself may be protective of inflammation and hyperglycemia.<sup>68</sup> |
| | Squalene and cholesterol biosynthesis | 280.974_191.9 | $[M-2H]^{2-}$ | Dimethylallyl diphosphate | DMAPP | Decreased | Dimethylallyl diphosphate is part of the mevalonate pathway and greater levels of squalene ultimately stimulates gluconeogenesis.<sup>69</sup> |
| HILIC | C21-steroid hormone biosynthesis and metabolism | 356.2627_23.3 | $[M-2H]^{2-}$ | 20α-Hydroxyprogesterone/Pregnenolone | 20α-OHP / Preg | Decreased | Positive association of progesterone with prediabetes and T2D, while pregnenolone was inversely associated with these conditions.<sup>70</sup> |
| | | 358.2773_24.3 | $[M-H_2O-H]^-$ | Allopregnanolone | AlloP | Decreased | Allopregnanolone is a neurohormone, and its signaling may mitigate oxidative stress and reduce 2-hydroxybutyrate which is associated with insulin resistance and hyperglycemia.<sup>71</sup> |
| | | 363.2207_240.7 | $[M-H_2O-H]^-$ | 11β,17α,21-Trihydroxypregnenolone | THP | Increased | This compound was negatively associated with hyperglycemia.<sup>70</sup> Therefore, pregnenolone signaling might be reduced in those with hyperglycemia. |
| | | 476.3725_23.7 | $[M-H_2O-H]^-$ | 5beta-cholestane-3alpha,7alpha,12alpha,25-tetrol | 5β-Cholestane-tetrol | Decreased | This compound is a precursor to cholic acid, a primary bile acid.<sup>72</sup> |
| | Glycosphingolipid metabolism | 322.042_74.9 | $[M-2H]^{2-}$ | CMP | - | Decreased | Intraperitoneal administration of CMP induced hyperglycemia in rats.<sup>73</sup> |
| | Leukotriene metabolism | 335.2192_71.5 | $[M-H_2O-H]^-$ | Leukotriene derivatives (e.g., 6,7-dihydro-5-oxo-leukotriene B4) | LT derivatives | Increased | Leukotriene metabolism is positively associated with poor glucose control in T1D.<sup>74</sup> |
| | | 336.2247_25.7 | $[M-H_2O-H]^-$ | Leukotrienes (e.g., 5-Hydroperoxy-6-trans-8,11,14-cis-eicosatetraenoate), Prostaglandins (e.g., Prostaglandin B1) | LTs, PGs | Increased | Leukotriene metabolism is positively associated with poor glucose control in T1D.<sup>74</sup> Both leukotriene and prostaglandin signaling are associated with inflammatory and stress signaling, upregulation after surgery may indicate upregulation of inflammation and tissue stress. |
| | | 351.2146_22.3 | $[M-H_2O-H]^-$ | | | Decreased | |
| Seventeen metabolites identified from group difference analysis showing statistically significant differences in temporal changes between pre-surgery baseline (T0) and 2 hours after surgery initiation (T1), distinguishing the stress hyperglycemia and non-hyperglycemia groups. Metabolites are organized by metabolic pathways identified through Mummichog. For each metabolite, the table provides chromatographic details (column type, m/z, retention time), adducts, annotations from Mummichog and KEGG, abbreviations, direction of change (increased/decreased), and clinical relevance with supporting references. |  |  |  |  |  |  |  |

### Metabolic Signature Predictive of Stress Hyperglycemia

Based on our previous findings, we developed machine learning models to predict SH using these metabolomic changes between T0 and T1 as input features. Model performance, evaluated via LOOCV, is summarized in **Supplementary Table 3**. The machine learning pipeline with logistic regression achieved the highest performance, with an AUC of 0.857, accuracy of 0.833, sensitivity of 0.893, and specificity of 0.750 (Figure 4C). Figure 4D displays the top 20 putative metabolites predictive of SH ranked by feature importance. One of these metabolites was among the 22 statistically significant metabolites identified in the whole cohort analysis, and three were among the 17 significant metabolites from the group difference analysis. This overlap shows convergence between statistical analyses and machine learning approaches in identifying key metabolic markers. Detailed m/z values, retention times, annotations, and abbreviations are provided in **Supplementary Table 9**.

### Biomarker Analysis and Multilevel Associations with Stress Hyperglycemia

Temporal trends for four biomarkers were assessed using the same NMEM approach applied to the whole cohort statistical analysis for metabolites: high-sensitivity C-reactive protein (hsCRP), soluble urokinase plasminogen activator receptor (SuPAR), brain-type natriuretic peptide (BNP), and high-sensitivity troponin (hsTnI). All biomarkers exhibited statistically significant temporal patterns, with all p-values less than 0.001. The estimated biomarker trajectories showed an overall increasing trend across the four time points, as shown in Figure 5A. Group difference statistical analysis was subsequently conducted to evaluate whether these temporal patterns differed by SH status. Statistical significance was observed for hsCRP (p<0.001) and SuPAR (p=0.014), while BNP (p=0.239) and hsTnI (p=0.065) showed non-significant trends. For hsCRP, participants with SH exhibited an increase between T0 and T1, while those without SH showed a delayed increase after T1, as shown in Figure 5B. For SuPAR, participants with SH had a steady increase from T0-T2 with a slight decrease at T3 (72-96 hours), whereas those without SH showed an upward trend after T1.

**Figure 5 A-D.**
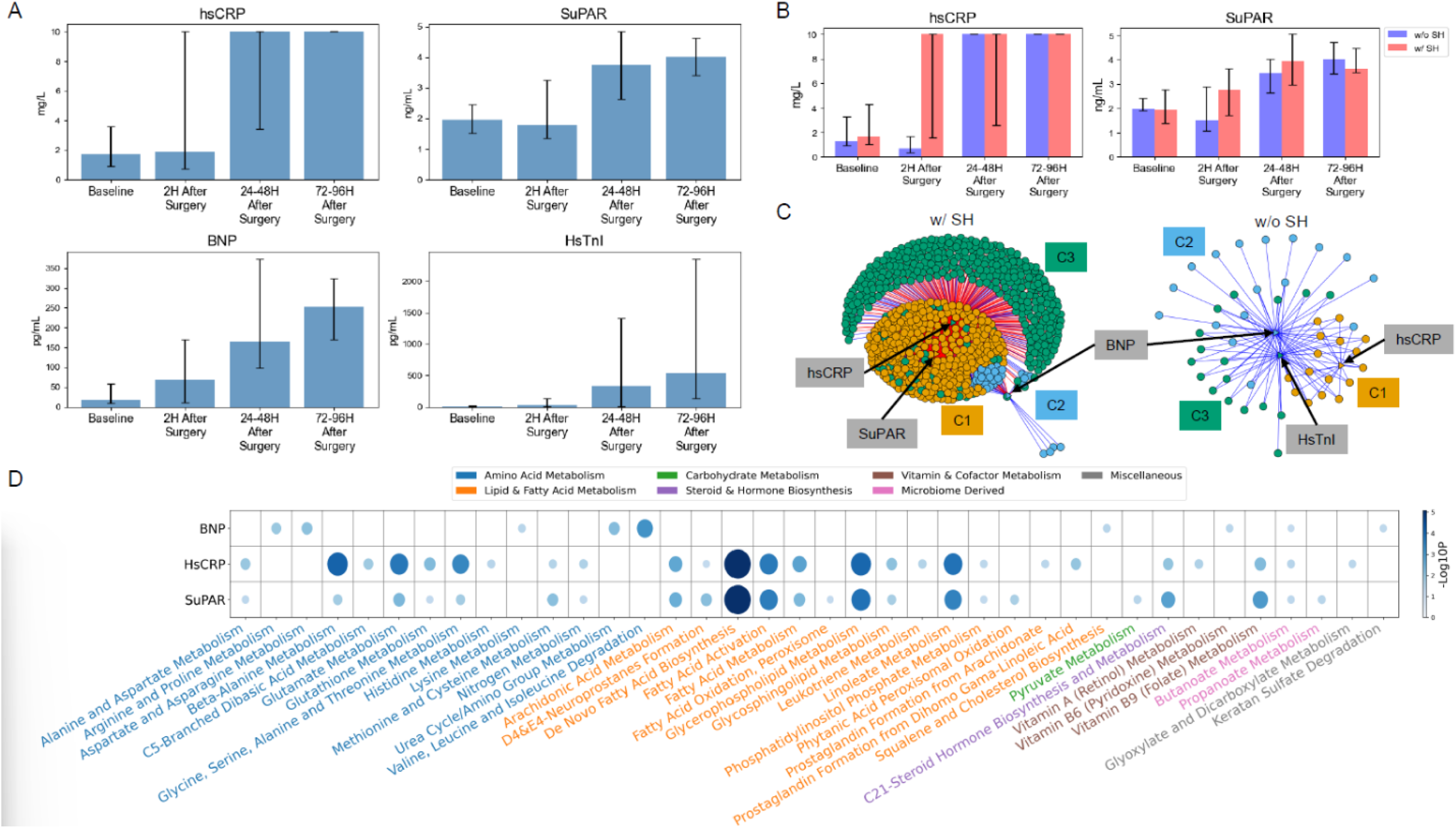
Biomarker trends during the perioperative period. **(A)** Biomarker trajectories across four perioperative time points from whole cohort analysis using nonlinear mixed-effects model. All biomarkers (hsCRP, SuPAR, BNP, hsTnI) demonstrated significant temporal variation (p < 0.001), with elevations during and after surgery. Data shown as median with interquartile range error bars. **(B)** Differential biomarker trajectories comparing participants with stress hyperglycemia (SH) (red bars) versus those without (blue bars). Significant differences were observed for hsCRP (p < 0.001) and SuPAR (p = 0.014), indicating distinct temporal patterns associated with hyperglycemia development. Data shown as median with interquartile range error bars. No difference was observed for BNP and hsTnI (not shown). **(C)** Network visualization of biomarker-metabolite associations stratified by hyperglycemia status. The left panel (w/ HGC) indicates dense metabolic networks showing SuPAR-associated community (C1, orange, 401 metabolites), BNP-associated community (C2, blue, 44 metabolites), and hsCRP-associated community (C3, green, 759 metabolites); right panel (w/o HGC) represents sparse networks with smaller communities for hsCRP (C1, orange, 17 metabolites), BNP (C2, blue, 17 metabolites), and hsTnI (C3, green, 40 metabolites). **(D)** Pathway enrichment analysis showing metabolic pathways most strongly correlated with each biomarker in participants with SH. Bubble size and color intensity indicate correlation strength (-log10 p-value, scale 0-5). BNP shows strongest associations with amino acid metabolism, while hsCRP and SuPAR are most associated with fatty acid metabolism

To explore biomarker-metabolite relationships, we mapped metabolomic communities in participants with and without SH. Participants with SH displayed substantially more metabolites associated with the inflammatory biomarkers hsCRP, SuPAR, and BNP compared to those without SH, as shown in Figure 5C. The strength of associations between each biomarker and metabolic pathways revealed distinct patterns between predominantly inflammatory biomarkers and BNP (Figure 5D). SuPAR showed the strongest associations with *de novo* fatty acid biosynthesis, folate, linoleate, and glycerophospholipids pathways. Amino acid metabolism (branched-chain amino acid degradation and the urea cycle) was strongly associated with BNP. *De novo* fatty acid biosynthesis, linoleate, amino acid metabolism (glycine, serine, alanine, threonine, glutamate, beta-alanine), and fatty acid activation pathway were most strongly associated with hsCRP.

## DISCUSSION

This longitudinal analysis revealed distinct and dynamic metabolic perturbations associated with surgical stress in a well-defined cohort without diabetes undergoing cardiac surgery. By integrating high-resolution metabolomics, continuous glucose monitoring (CGM), and biomarker data, we identified coordinated changes across biochemical pathways linked to inflammation, lipid remodeling, amino acid metabolism, and bile acid signaling, changes that were more pronounced in individuals who developed SH. These findings provide mechanistic insights into the metabolic underpinnings of surgical stress and highlight potential biomarkers that may inform perioperative risk stratification and potential targets for future therapeutic strategies.

### Baseline Pathway Differences and Predisposition to Stress Hyperglycemia

Several metabolic pathways were different at baseline in those who developed SH, including Carnitine shuttle, Branched-chain amino acid (BCAA) metabolism, Bile acid biosynthesis, Inflammatory signaling (arachidonic acid metabolism), Microbiome-related metabolism (propanoate metabolism). Many of these pathways have been previously linked to diabetes risk. Their presence preoperatively suggests that individuals predisposed to SH may have underlying metabolic profiles favoring dysglycemia, even before surgical stress. The clinical implications of these preoperative signatures, particularly in patients without diagnosed diabetes, warrant further study in the perioperative and post-discharge periods.

### Perioperative Dynamics and Dominant Temporal Patterns

Across temporal analyses (whole cohort, CGM-integrated, and SH-stratified), pathways not evident at baseline, most prominently glycerophospholipid and glycosphingolipid metabolism, emerged after including perioperative timepoints. A characteristic “increased-during-surgery” profile was observed, marked by a sharp rise at T1 followed by partial resolution at T2–T3 Complementary clusters exhibited a “decreased-during-surgery” pattern, with T1 nadirs and rebound thereafter. These coordinated lipid trajectories mirror the perioperative glucose dynamics and are biologically consistent with membrane remodeling and tissue injury/repair. Notably, the amplitude and persistence of these changes were greater in participants who developed SH, with slower normalization and higher trajectories across time. Together, the magnitude and synchrony of these lipid-centered shifts suggest that a subset of patients is more metabolically vulnerable to stress-induced derangements during cardiac surgery.

### Key Metabolites and Pathway-Specific Insights

#### Lipid-Based Inflammatory Signaling

Multiple lipid-derived inflammatory pathways were dominant postoperatively. Arachidonic acid metabolism, prostaglandin formation, and leukotriene metabolism were upregulated in SH, consistent with prior evidence linking them to inflammation, insulin resistance, and poor glucose control.^3–6^ Sphinganine and sphinganine-1-phosphate, precursors in ceramide biosynthesis, were associated with SH, suggesting ceramide-mediated metabolic disruption as a potential mechanism for perioperative insulin resistance.^7,8^ Free fatty acids (FFAs) may further exacerbate this state by stimulating hepatic glucose production, increasing oxidative stress, and preventing insulin-mediated suppression of glycogenolysis.^9^

#### Amino Acid and Sulfur Pathways

Methionine and cysteine metabolism, closely linked to homocysteine regulation, was altered. Elevated homocysteine is a cardiovascular risk factor, and changes post-CABG may reflect altered folate-dependent methylation.^10^ Glycine, alanine, and gluconeogenic amino acids may reflect enhanced hepatic glucose output.^11^ increased release of glucose from the liver has been shown to occur with elevated gluconeogenic amino acids, suggesting one way in which altered amino acid metabolism can lead to SH.^12^ In addition, alanine is a well-known substrate of gluconeogenesis, suggesting that elevated alanine levels may upregulate gluconeogenesis, leading to more glucose production and subsequently greater glucose levels.^12^

#### Bile Acid Signaling

Bile acids were associated with SH. While bile acids can have protective metabolic effects via FXR and TGR5 signaling, higher levels in SH may reflect glucose-driven synthesis rather than a protective adaptation.^13,14^

#### Histidine Metabolism

Lower levels of 4-imidazolone-5-propanoate in SH suggest increased conversion of histidine to imidazole propionate, a metabolite linked to impaired glucose metabolism.^15^

### Biomarker–Metabolite Associations

To integrate biochemical changes with systemic signals, we examined correlations between key biomarkers and metabolites: SuPAR was associated with de novo fatty acid biosynthesis, glycerophospholipid metabolism, and linoleate, consistent with inflammation and immune activation. SuPAR is linked to diabetes, cardiovascular disease, and adverse renal outcomes.^16,17^ BNP was linked to alterations in amino acid metabolism, particularly branched-chain amino acid degradation and urea cycle pathways, suggesting possible nitrogen handling dysregulation in the context of CABG and CAD. hsCRP had associations similar to SuPAR, reinforcing the central role of inflammation in SH pathophysiology. These biomarkers may serve as accessible surrogates for deeper metabolic changes and help stratify perioperative risk.

### Interpretation in Context

The convergence of lipid-derived inflammatory pathways, amino acid dysregulation, and bile acid signaling highlights the multifactorial metabolic stress response to surgery. In SH, this response is amplified, possibly reflecting a combination of: a) pre-existing metabolic susceptibility (evident at baseline), b) surgical tissue injury and membrane remodeling, and c) inflammatory-hormonal activation (cortisol, catecholamines, cytokines). This metabolic fingerprint may also have implications for post-discharge diabetes risk assessments in susceptible individuals.

### Strengths and Limitations

This study integrates metabolomics, continuous glucose monitoring, and biomarker data in a non-diabetic surgical cohort, offering a multidimensional view of metabolic responses to surgical stress. A multi-timepoint design enabled dynamic trajectory analysis, while advanced analytical methods captured coordinated pathway-level changes beyond individual metabolites.

Limitations include a relatively small sample size, which reduced statistical power—particularly for subgroup and pathway-level analyses—and may limit generalizability. Although CGM was calibrated to point-of-care testing, perioperative factors such as temperature, perfusion, and hemodynamic instability may have affected accuracy. Residual confounding from hormonal fluctuations, genetic variability, and differences in surgical severity cannot be excluded; notably, non-SH group surgeries may have been less severe than those in the SH group. Replicating the complexity of human surgical stress in controlled models also remains challenging. Finally, pathway enrichment analyses were exploratory, intended to generate mechanistic hypotheses; confirmation in larger, independent cohorts is needed to establish clinical relevance.

### Conclusion

Patients who develop SH after CABG exhibit distinct baseline metabolic signatures, encompassing carnitine shuttle activity, branched-chain amino acid metabolism, bile acid biosynthesis, inflammatory processes, and microbiome-related pathways, that persist or intensify after surgery. Surgical stress further unmasks additional perturbations, particularly in glycerophospholipid and glycosphingolipid metabolism, which may reflect tissue injury. Integration of machine learning with pathway analysis offers a potential framework for precision medicine strategies to predict and mitigate SH. Given the close links between these metabolic stress signatures, inflammation, and insulin resistance, they may also serve as early indicators of postoperative diabetes risk, underscoring the need for validation in larger, independent cohorts.

## METHODS

### Study Population

Participants comprised patients without diabetes (blood glucose < 126 mg/dL and HbA1c < 6.5%) undergoing CABG surgery at three academic hospitals (Emory University Hospital, Emory University Hospital Midtown, Grady Memorial Hospital) in Atlanta, Georgia. The study was approved by the Institutional Review Board at Emory University (Protocol IRB00097963). All participants provided written informed consent prior to enrollment.

Participants in this study were part of a nested case-control study linked to a clinical trial testing a GLP-1 receptor agonist in the perioperative period (NCT03743025). We collected the following variables during this study: 1) demographic characteristics (age, sex [male, female], race-ethnicity [White, Black, Hispanic, Others]); 2) clinical information (weight, BMI, smoking status [never smoked], cardiac ejection fraction, HbA1c, blood glucose at consent, APACHE II score, length of stay); 3) surgical information (type of surgery [open, robotic], cardiopulmonary bypass); 4) history of cardiovascular or other diseases (previous cardiac condition, hyperlipidemia, hypertension, infection); 5) medication exposure (vasopressors or inotropes, norepinephrine, vasopressin, dobutamine, milrinone).

### Data Collection and Variable Specification

Blood samples were collected at four perioperative time points from T0 to T3. Participants who received dulaglutide before surgery (eight) and those who received steroids during surgery (six) were excluded. Statistical comparisons between groups were performed using the two-sample t-test for continuous variables and chi-squared tests for categorical variables. All data was securely stored in REDCap.

#### Point-of-Care Testing

Accu-Chek Inform II glucometer (Emory University Hospital and Emory University Hospital Midtown) and Nova StatStrip (Grady Memorial Hospital) were used to measure glucose from capillary or arterial blood. Point-of-Care (POC) glucose values were checked during the perioperative period, both during and after surgery, based on hospital protocols or anesthesiology discretion.

#### Continuous Glucose Monitoring

Factory-calibrated, blinded Dexcom G6-pro CGM devices were placed on participants prior to CABG surgery. The device measures glucose every five minutes for up to 10 days. The device was previously shown to have an overall mean absolute relative difference (MARD) of 12.8% relative to POC capillary glucose previously in inpatient settings.^18^

#### Biomarkers

Plasma was collected to measure biomarkers. hsCRP was measured by FirstMark and suPAR was measured using suPARnostic kit from Virogates, Copenhagen, Denmark.

Trends between biomarkers, metabolites, and SH were assessed by metabolome-wide association studies (xMWAS) and integrative network analysis.

#### Metabolomics

Blood samples were collected at four time points, and high-resolution untargeted metabolomics analysis was performed at the Clinical Biomarkers Laboratory using established protocols with C18 under negative ionization mode (8,631 mass spectrometry features) and HILIC under positive ionization mode (10,919 mass spectrometry features).^19–21^ For data pre-processing, metabolites with zero values were imputed using half the minimum detected value for that metabolite. Metabolite intensities were log-transformed prior to statistical analysis.

### Stress Hyperglycemia Definition

SH was defined as meeting either of the following criteria during the period from 4 to 28 hours after the start of surgery (24-hour duration): 1) three or more POCT measurements above 140 mg/dL, or 2) any single POCT measurement above 180 mg/dL. Perioperative insulin administration followed standard institutional protocols.

### Statistical Analysis

#### Baseline Group Difference Analysis

Baseline differences in metabolite levels between groups were assessed using two-sample t-tests. For each metabolite, a separate test was performed using baseline (T0) values to compare group means, where a significant p-value indicated that the metabolite showed a statistically significant difference at baseline.

#### Whole Cohort Analysis

Temporal changes in metabolite levels were investigated using NMEM.^22^ For each metabolite, two alternative models were compared to determine whether the metabolite exhibited significant temporal variation (full model) or remained stable over time (null model). Specifically, the full model took the following form:

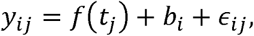

where *y_ij_* denotes the metabolite level for participant *i* at time point *j*, *f*(*t_j_*) is a non-linear function of time modeled using natural splines with three degrees of freedom (excluding the intercept), 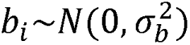 is a participant-specific random intercept, and 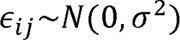 is the residual error. In contrast, the null model assumes no temporal change of the metabolite by replacing *f*(*t_j_*) with a constant intercept.*β_0_* The full and null models were compared using likelihood ratio tests, where a significant p-value indicated that the metabolite exhibited a statistically significant temporal change.

#### Group or Clinical Difference Analysis

To assess whether temporal trajectories of metabolite levels were modified by a variable of interest *Z_i_* (binary group or continuous clinical measure), a NMEM was extended to include an interaction between time and *Z_i_*. The full model takes the following form:

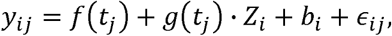

where *g(t_j_)* is a non-linear function of time modeled using natural splines and 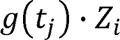 captures the interaction, allowing the time trajectory to vary by *Z_i_*. In contrast, the null model is excluded in this interaction. Likelihood ratio tests were used to compare models, with significant p-values indicating that temporal patterns differed by *Z_i_*.

#### Pathway Enrichment Analysis

To identify biologically meaningful pathways associated with temporal changes in metabolite levels, pathway enrichment analysis was performed using Mummichog version 2.7.0.^23,24^ Separate analyses were conducted for C18 and HILIC datasets, with metabolites analyzed alongside their corresponding p-values derived from NMEM. The p-value cutoff was set to 0.05, and 1,000 permutations were used to assess statistical significance. The identified significant pathways provided insights into the underlying metabolic processes associated with the observed temporal metabolite dynamics.

To uncover common temporal trends among metabolites within significant biological pathways, we analyzed the subset of metabolites included in significant pathways identified through pathway enrichment analysis. Metabolites from both C18 and HILIC datasets were combined, and temporal trajectories were derived from NMEM-fitted values using the full model. These trajectories were clustered using the DBSCAN algorithm,^25^ with similarity defined as one minus the Pearson correlation coefficient between metabolite trajectories. The resulting clusters revealed dominant temporal patterns, reflecting groups of metabolites that exhibited similar dynamics over time.

### Machine Learning Pipeline

A participant-level classifier was developed to predict SH development using metabolomic data. The initial feature set comprised empirical metabolites identified through Mummichog pathway enrichment analysis: 1,455 putative metabolites from C18 and 1,320 from HILIC (2,775 total metabolites). Model performance was evaluated using leave-one-out cross-validation (LOOCV) based on area under the receiver operating characteristic (ROC) curve (AUC), accuracy, sensitivity, and specificity.

For each LOOCV training set, recursive feature elimination (RFE) was employed,^26–29^ which iteratively removed 10% of features with the lowest RFE importance scores until 25 features remained. RFE importance scores were determined using ridge logistic regression coefficients to avoid overfitting. Partial least squares (PLS) regression was subsequently applied to derive a single composite feature, with age, sex, and BMI included as additional covariates, resulting in four predictors. Four classification algorithms were then evaluated based on these predictors: Ridge logistic regression, support vector machine (SVM) classifier, Random Forest, and Multilayer perceptron (MLP) classifier.

Feature importance for interpretation was calculated within the LOOCV framework. In each iteration, logistic coefficients were multiplied by the corresponding PLS coefficients to account for magnitude and direction. The absolute values of these combined coefficients were averaged across all LOOCV iterations, and the top 20 metabolites were selected based on these averaged absolute importance values.

### Integrative Network Analysis of Clinical and Metabolic Data

To investigate metabolic pathways associated with SH development, data-driven integrative analysis was performed using xMWAS.^30^ Eight clinical covariates at T0 and T1 (age, sex, BMI, SuPAR, BNP, hsTnI, hsCRP, and SH status) were integrated with corresponding metabolic profiles using PLS canonical regression. A multilevel community detection algorithm identified connected groups of variables that exhibited strong intra-community connections while maintaining sparse connections with the broader network. Network edges were retained based on threshold criteria of the absolute correlation coefficient greater than 0.6 and p-value less than 0.05 as determined by Student’s t-test.

### Manual Annotation of Metabolites

For metabolites not annotated by Mummichog, manual annotation was performed using the Human Metabolome Database (HMDB) version 5. m/z ratios were queried using the liquid chromatography-mass spectrometry (LC-MS) search feature with separate searches for positive and negative ionization modes. For positive mode, the following adducts were prioritized: *[M+H]^+^*, *[M+ACN+H]^+^*, *[M+ACN+Na]^+^*, with additionally consideration of *[M+NH_4_]^+^*, *[M+Na]^+^*, *[M+H−H_2_O]^+^*, *[M+K]^+^*, and *[M+H+Na]^+^*. For negative mode, *[M−H]^−^*, *[M−H_2_O−H]^−^*, and *[M+Cl]^−^*adducts were included. All searches employed a five-ppm mass tolerance between measured and theoretical m/z values. Annotation assignment followed a hierarchical prioritization scheme. First, molecules not endogenously produced by human or microbiome metabolism were given lower priority to focus on biologically relevant metabolites. Second, annotations with the smallest mass deviation (lowest delta m/z) received highest priority. Lastly, biological relevance and isotopic consistency of the same chemical formula were used to assign priority at the discretion of the researchers.

### CGM Imputation and Calibration

To address the challenges of missing or inaccurate values in CGM data during the perioperative period,^31^ we developed an imputation and calibration protocol that integrates CGM and POCT measurements, which includes three steps.

#### Step 1: Time Alignment

Prior studies have reported a temporal lag between POCT and CGM, with POCT measurements typically preceding CGM readings.^32–34^ To account for this lag, Pearson correlations between CGM and POCT were calculated using time shifts ranging from 5 to 25 minutes. The time shift with the highest correlation was selected as the optimal lag for each participant. The CGM and POCT time series were then aligned based on this optimal lag.

#### Step 2: Imputation

Unlike conventional imputation methods that rely solely on observed CGM values to estimate missing data, our approach leveraged the availability of paired POCT and CGM measurements to improve imputation quality. Specifically, for time points where both CGM and POCT readings are available, we modeled their relationship using local linear regression and used this model to predict missing CGM values from corresponding POCT measurements. This enabled imputation based on a clinically verified glucose reference rather than simply relying on internal CGM trends. Specifically, at each time point *t* with a missing CGM value and a corresponding POCT, a local window *W_t_* was defined, which consisted of nine nearby time points with available CGM-POCT pairs. Within this window, the following local regression was fitted:

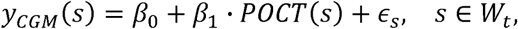

where *Y_CGM_*(*S*) and *POCT*(s) denote the CGM and POCT values at time *s*, respectively, and 𝛜*_s_* is the error term. This model allowed for the local and non-stationary relationship between CGM and POCT to be captured. To estimate *β_0_* and *β_1,_* we proposed to solve a constrained optimization problem as follows:

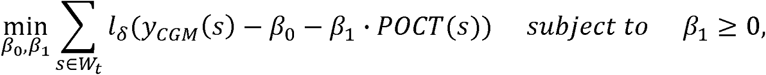

where *l_δ_* (․) is the Huber loss function between the true and predicted CGM values.^35^ This loss function was chosen for its robustness to outliers, helping to mitigate the impact of occasional noise or unreliability in CGM readings. The non-negativity constraint on *β_0_* reflects the expected positive association between CGM and POCT values. After solving the optimization, the estimated 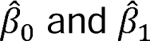 were used to impute the missing CGM value at time *t* using the corresponding POCT value. For remaining CGM time points without paired POCT data, linear interpolation between neighboring CGM values was used to complete the time series.^36–38^

#### Step 3: Calibration

To further improve the accuracy of CGM data, we developed a calibration step to account for time-varying sensor behavior, such as signal drift or adaptation effects.^39,40^ For this, a separate local linear regression model was fitted to predict POCT values using CGM readings (both observed and imputed from Step 2) and the time since sensor insertion.

Specifically, the following regression was fitted:

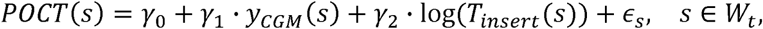

where *T_insert_*(*S*) denotes the time since sensor insertion at time *S* and the logarithmic transformation was used to capture non-linear behavior related to sensor adaptation over time. To estimate *γ_0_*, *γ_1_* and *γ_2_* we proposed to solve a similar constrained optimization as in Step 2 as shown below:

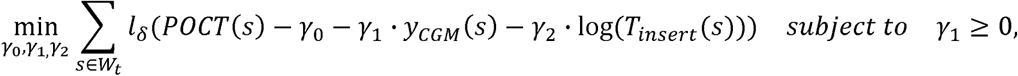

The estimated 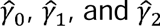 were utilized to predict the POCT values, which served as calibrated CGM readings. This method adjusts CGM readings by accounting for both their instantaneous relationship with POCT and time-dependent sensor drift, enhancing calibration accuracy.

## Author contributions

### Contributions

D.K. and J.B. contributed equally to this work. D.K.,J.B. drafted the initial manuscript. D.K. conducted pathway, AI/ML, and CGM analyses. J.B., JF, L.G., A.A., S.K., N.G., S.U., H.H., interpretated metabolites and pathway relevance, C.P.G. enrolled participants and collected samples, AC and M.H. participated in recruitment and clinical care. RGK, TZ, DPJ, and AAQ provided feedback on study design reviewed the manuscript. MRS led and supervised metabolomics analyses, J.L., led and supervised pathway, AI/ML, and CGM analyses, F.J.P. secured funding, and supervised the overall project. All authors: review & editing

