## Supplementary Information for "Metabolic Trajectories During Surgical Stress in Patients Undergoing Cardiac Surgery"

**Funding:** The authors acknowledge the support from the National Institute of General Medical Sciences Award number K23GM128221 (FJP), the National Institute of Environmental Health Sciences Award number P30 ES019776 (DPJ), the Veterans Health Administration Career Development Award Number (5IK2BX005913-02; MRS), the Clinical Biomarkers Laboratory, and an ancillary grant from Dexcom. JF is supported in part by the National Center for Advancing Translational Sciences of the National Institutes of Health under award number UL1 TR002378 and TL1TR002382.

**Corresponding Authors**:

Francisco J Pasquel, MD, MPH

Associate Professor

Department of Medicine

Division of Endocrinology

Emory University

Jing Li, PhD

Professor

H. Milton Stewart School of Industrial and Systems Engineering

Georgia Institute of Technology

**CGM Calibration**

Accurate CGM calibration is critical for reliable estimation of glycemic variability and time above range. To validate our imputation and calibration method, we analyzed data from 50 participants who had both CGM and point-of-care testing measurements. On average, each participant had 31.36 POCT measurements (SD=11.53) and 1,620.48 CGM readings (SD=597.88). An average of 43.2 CGM readings (SD=52.27) per participant were missing and required imputation prior to calibration. We employed leave-one-out cross validation to train and evaluate our proposed imputation and calibration method. To isolate the contribution of each methodological component, we compared our approach against three alternative methods: (1) Huber Regression: similar to our proposed method but without the non-negativity constraint on the regression coefficient; (2) Linear Regression: a standard local regression approach using least squares loss instead of Huber loss, without constraint. (3) Uncalibrated CGM: raw CGM data without any calibration step.

Performance was evaluated using two standard metrics: Mean Absolute Difference (MAD) and Mean Absolute Relative Difference (MARD) between calibrated CGM predictions and reference POCT values. Results for each method are summarized in **Table S1.** Our proposed method achieved superior performance with the lowest error rates (MAD=11.68, MARD=9.40%) and the smallest standard deviations across both metrics. **Figure S1** shows four examples of calibrated CGM using the proposed method, overlaid with original CGM readings and POCT measurements.

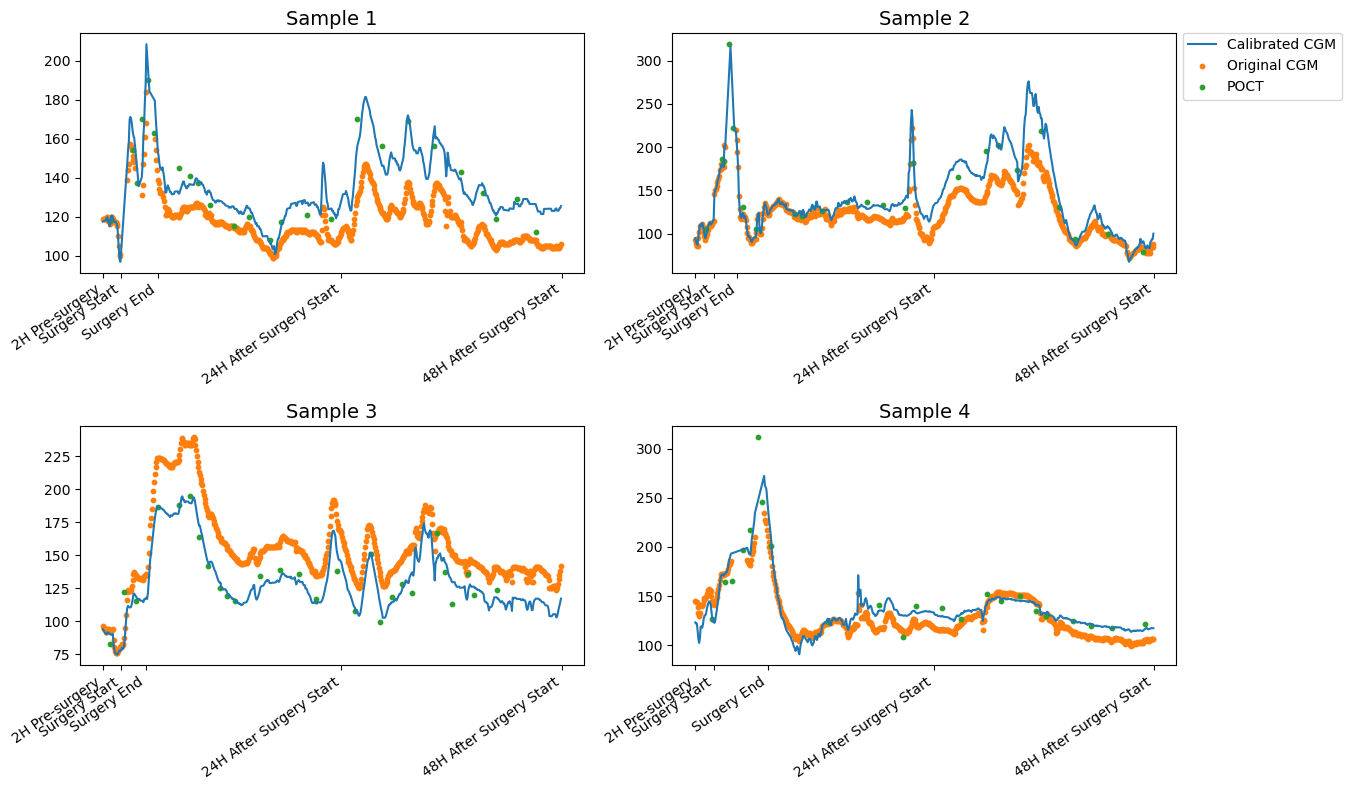
**Supplementary Table 10.** Performance Comparison of CGM Calibration Methods

|  | Proposed Method | Huber Regression | Linear Regression | Uncalibrated CGM |
| --- | --- | --- | --- | --- |
| MAD | 11.68 (3.92) | 11.91 (4.05) | 12.40 (4.44) | 33.41 (21.09) |
| MARD | 9.40% (3.26) | 9.68% (3.54) | 10.10% (3.91) | 25.63% (15.85) |
| Comparative evaluation of CGM calibration approaches using leave-one-out cross-validation. Performance was assess using Mean Absolute Difference (MAD) in mg/dL and Mean Absolute Relative Difference (MARD) as percentage between calibrated CGM predictions and reference POCT values, with standard deviations in parenthesis. | | | | |

**Supplementary Figure 1.** Comparison of calibrated CGM (blue line), original CGM readings (orange dots), and POCT measurements (green dots) across four participants during the perioperative period. The proposed calibration method adjusts the original CGM readings to better align with POCT reference values.

**Supplementary Table 1.** Participants Characteristics

|  | **Stress Hyperglycemia**  **(n=28)** | **No Stress Hyperglycemia (n=20)** | **p-value** |
| --- | --- | --- | --- |
| Age | 62.6 (11.9) | 65.8 (6.5) | 0.24 |
| Sex  Male | 26 (93%) | 16 (80%) | 0.38 |
| Race/ethnicity  White  Black  Hispanic  Other | 16 (57%) 7 (25%) 2 (7%) 3 (11%) | 13 (65%) 5 (25%)  0 (0%) 2 (10%) | 0.67 |
| Weight (kg) | 91.7 (17.2) | 85.2 (16.0) | 0.19 |
| BMI (kg/m^2^) | 28.9 (4.3) | 27.5 (4.8) | 0.30 |
| Smoking Status (Never Smoked) | 9 (32%) | 10 (50%) | 0.34 |
| Cardiac Ejection Fraction (EF)% | 52.7 (9.7) | 54.3 (9.9) | 0.58 |
| HbA1c (%) | 5.6 (0.4) | 5.6 (0.3) | 0.66 |
| Blood Glucose at Consent | 99.7 (12.9) | 96.5 (14.8) | 0.43 |
| APACHE II Score | 14.9 (6.1) | 13.4 (5.0) | 0.36 |
| Length of Stay (days) | 7.6 (8.3) | 5.2 (1.5) | 0.14 |
| Type of Surgery  Open  Robotic | 27 (96%) 1 (4%) | 18 (90%) 2 (10%) | 0.76 |
| Cardiopulmonary Bypass | 25 (89%) | 11 (55%) | 0.02 |
| Previous Cardiac Condition | 22 (79%) | 16 (80%) | 1.00 |
| Hyperlipidemia | 24 (86%) | 14 (70%) | 0.34 |
| Hypertension | 24 (86%) | 18 (90%) | 1.00 |
| Infection | 1 (4%) | 1 (5%) | 1.00 |
| Vasopressors or Inotropes | 26 (93%) | 20 (100%) | 0.63 |
| Norepinephrine | 24 (92%) | 19 (95%) | 1.00 |
| Vasopressin | 7 (28%) | Yes: 2 (10%) | 0.26 |
| Dobutamine | 4 (16%) | Yes: 4 (20%) | 1.00 |
| Milrinone | 7 (27%) | Yes: 1 (5%) | 0.12 |
| Demographic Characteristics of participants stratified stress hyperglycemia. Continuous variables are presented as mean (standard deviation) and categorical variables as count (percentage). Two participants in the hyperglycemia group had missing data each for norepinephrine and milrinone administration; three participants each for vasopressin and dobutamine administration. Statistical significances were calculated using two-sample t-test for continuous variables and chi-squared tests for categorical variables. | | | |

**Supplementary Table 2**

**Supplementary Table 3.** Predictive Performance Comparison for Stress Hyperglycemia

| **Baseline Analysis** | | | | |
| --- | --- | --- | --- | --- |
|  | **AUC** | **ACC** | **Sensitivity** | **Specificity** |
| Logistic Regression | 0.7929 | 0.7500 | 0.7857 | 0.7000 |
| SVM | 0.7911 | 0.7500 | 0.7857 | 0.7000 |
| Random Forest | 0.7708 | 0.7708 | 0.8929 | 0.6000 |
| MLP | 0.7292 | 0.7292 | 0.7500 | 0.7000 |
| **Change from Baseline to T1** | | | | |
|  | **AUC** | **Accuracy** | **Sensitivity** | **Specificity** |
| Ridge Logistic Regression | 0.857 | 0.833 | 0.893 | 0.750 |
| SVM Classifier | 0.743 | 0.813 | 0.893 | 0.700 |
| Random Forest | 0.823 | 0.813 | 0.786 | 0.850 |
| MLP Classifier | 0.850 | 0.792 | 0.714 | 0.900 |
| Performance metrics of machine learning models for predicting stress hyperglycemia using metabolomic data from baseline and changes between T0 and T1. Evaluation metrics include area under the curve (AUC), accuracy, sensitivity, and specificity, assessed using leave-one-out cross-validation. | | | | |

**Supplementary Table 4.** Top 20 Metabolites Ranked by Feature Importance for Baseline Prediction of SH.

| **Rank** | **Column type** | **m/z_RT (s)** | **Adducts** | **Mummichog, KEGG Label** | **Abbreviation** | **Relevance** |
| --- | --- | --- | --- | --- | --- | --- |
| 1 | HILIC | 252.1786_25.1 | M(C13)+H[1+] | 3-Hydroxylidocaine | 3-OH-Lidocaine-C13 | No relevance to hyperglycemia or cardiac outcomes found. |
| 2 | C18 | 239.0155_23.9 | M-H[-] | L-Cystine; L-Dicysteine; L-alpha-Diamino-beta-dithiolactic acid | Cys/CySS/DithLA | In rats with diabetes, L-cysteine (L-cystine is a derivative of L-cysteine) supplementation is associated with decreased glucose values and insulin resistance.^1^ |
| 3 | HILIC | 283.2267_237.8 | M-CO+H[1+] | *Monoacylglycerol | MG | Monoacylglycerols were found to be higher in db/db mice which had increased intestinal inflammation.^2^ |
| 4 | HILIC | 311.2569_20.3 | M-CO2+H[1+] | Trioxilin A3 | - | Trioxilin A3 is involved in insulin production and inflammatory responses.^3^ |
| 5 | HILIC | 142.0693_62.5 | M(C13)+H[1+] | 3-Methylimidazoleacetic acid | 3-MIAA | See table 9. |
| 6 | HILIC | 146.0925_219.6 | M+H[1+] | 4-Guanidinobutanoate | 4-GBA | No direct evidence of ties to hyperglycemia or cardiovascular outcomes. However, 4-guanidinobutanoate is involved in inflammatory pathways.^4^ |
| 7 | HILIC | 251.1752_25.0 | M+H[1+] | 3-Hydroxylidocaine | 3-OH-Lidocaine | No relevance to hyperglycemia or cardiac outcomes found. |
| 8 | HILIC | 322.8029_159.1 | M+HCOONa[1+] | Iodine; I2 | Iodine | Increased iodine levels have been associated with type 2 diabetes.^5^  Atrial fibrillation can also result from hyperthyroidism from high iodine intake.^6^ |
| 9 | HILIC | 263.1114_65.1 | M+Na[1+] | beta-Alanyl-N(pi)-methyl-L-histidine (anserine) | Anserine | Anserine may prevent cardiac dysfunction.^7^ |
| 10 | C18 | 112.004_292.1 | M-H2O-H[-] | Iminoaspartate | IminoAsp | No relevance to hyperglycemia or cardiac outcomes found. |
| 11 | C18 | 345.2068_173.4 | M-H[-] | 11-Deoxycortisol; Cortodoxone | 11-Deoxycortisol | No direct evidentiary relation.  However, 11-deoxycortisol is a precursor to cortisol, a glucocorticoid with gluconeogenic effects potentially related to hyperglycemia. |
| 12 | HILIC | 432.3105_27.8 | M-H2O+H[1+] | Chenodeoxyglycocholate$ | CDGCA | No direct evidentiary relation.  However, chenodeoxycholate is a bile acid, a conjugate of chenodeoxycholate and glycine. Chenodeoxycholate is a primary bile acid with direct glucose and metabolic signaling roles through FXR and TGR5. |
| 13 | HILIC | 173.0921_62.1 | M+Na[1+] | Perillyl aldehyde; Perillaldehyde | PerAld | Exogenous plant essential oil extract. |
| 14 | C18 | 464.3014_24.4 | M-H[-] | glycocholate | GCA | Glycocholate, a conjugate of the primary bile acid cholic acid and glycine, is associated with gestational diabetes via hepatic steatosis.^8^ |
| 15 | C18 | 437.2927_223.2 | M+HCOO[-] | Deoxycholic acid; Deoxycholate | DCA | See table 9. |
| 16 | HILIC | 86.06_32.1 | M-CO2+H[1+] | 5-Oxoproline | 5-OxoPro | A metabolite of the glutathione synthesis and degradation pathways, 5-oxoproline is associated with myocardial dysfunction and oxidative stress^9^ and is higher in those with type 2 diabetes.^10^ |
| 17 | C18 | 315.198_142.7 | M-H2O-H[-] | Prostaglandin A2; PGA2; Medullin | LTs, PGs | See table 9 |
| 18 | HILIC | 238.092_109.0 | M+H[1+] | fructoseglycine | FruGly | Fructoseglycine is formed by non-enzymatic glycation of a fructose to glycine, can be considered an advanced glycation end product (AGE) which may be reflective of short-term glycemia similar to a fructosamine measurement or HbA1c. |
| 19 | C18 | 130.0509_19.2 | M+ACN-H[-] | D-Lactate | - | See tables 6,7. |
| 20 | C18 | 109.0409_159.6 | M-H[-] | Imidazole-4-acetaldehyde; Imidazole acetaldehyde | Im4Ald | Formed from the deamination breakdown of histamine, therefore is related to histidine metabolism. May be related to surgical outcomes as it was found in an untargeted metabolomics by LC-MS to be different between surgical patients with higher and lower opioid use after surgery.^11^ |
| This table lists the top 20 metabolites for predicting SH using ridge logistic regression. Rank indicates the importance of each metabolite, with rank 1 being the most important. Chromatography details are outlined by column type, m/z ratio, and retention time. Mummichog and KEGG provided chemical annotations and metabolite abbreviations are provided. Relevance indicates the metabolite’s relevance to the outcomes of interest. When the relevance is already indicated in other tables, the table is referenced. *Metabolite identified by manual annotation. | | | | | | |

**Supplementary Table 5.** Pathways Associated with Glycemic Variability (Coefficient of Variation)

| **Column** | **Pathway** | **Overlap size /**  **Pathway size** | **p-value** |
| --- | --- | --- | --- |
| C18 | Glycosphingolipid metabolism | 11 / 21 | 0.016 |
|  | Glycerophospholipid metabolism | 19 / 43 | 0.018 |
|  | Purine metabolism | 23 / 55 | 0.021 |
|  | Vitamin B9 (folate) metabolism | 4 / 5 | 0.023 |
|  | Glycine, serine, alanine and threonine metabolism | 15 / 33 | 0.026 |
|  | Beta-Alanine metabolism | 7 / 13 | 0.042 |
|  | Alanine and Aspartate Metabolism | 12 / 27 | 0.047 |
|  | Propanoate metabolism | 4 / 6 | 0.049 |
| HILIC | Glycosphingolipid biosynthesis - globoseries | 4 / 5 | 0.005 |
|  | Glycosphingolipid metabolism | 10 / 22 | 0.005 |
|  | Glycosphingolipid biosynthesis - ganglioseries | 5 / 8 | 0.007 |
|  | Phosphatidylinositol phosphate metabolism | 9 / 22 | 0.013 |
|  | Prostaglandin formation from arachidonate | 9 / 22 | 0.013 |
|  | Glycosphingolipid biosynthesis - lactoseries | 3 / 4 | 0.016 |
|  | Blood Group Biosynthesis | 3 / 4 | 0.016 |
|  | Glycosphingolipid biosynthesis - neolactoseries | 3 / 4 | 0.016 |
|  | Keratan sulfate biosynthesis | 3 / 4 | 0.016 |
|  | O-Glycan biosynthesis | 3 / 4 | 0.016 |
|  | Leukotriene metabolism | 9 / 23 | 0.017 |
|  | Sialic acid metabolism | 6 / 16 | 0.049 |
| Metabolic pathways significantly associated with coefficient of variation from continuous glucose monitoring during the first 48 hours of surgery. Pathway enrichment analysis was performed using Mummichog for C18 and HILIC chromatography. Pathway size refers to the total number of putative compounds associated with a given pathway, while overlap size represents the number of significant putative compounds detected. P-values indicate statistical significance of pathway enrichment. | | | |

**Supplementary Table 6.** Pathways Associated with Time Spent Above 140mg/dL

| **Column** | **Pathway** | **Overlap size /**  **Pathway size** | **p-value** |
| --- | --- | --- | --- |
| C18 | Pyrimidine metabolism | 24 / 60 | 0.001 |
|  | Pentose phosphate pathway | 10 / 19 | 0.003 |
|  | Pentose and Glucuronate Interconversions | 7 / 11 | 0.003 |
|  | Vitamin B9 (folate) metabolism | 4 / 5 | 0.009 |
|  | Aspartate and asparagine metabolism | 17 / 47 | 0.015 |
|  | Propanoate metabolism | 4 / 6 | 0.019 |
|  | Selenoamino acid metabolism | 5 / 9 | 0.024 |
|  | Chondroitin sulfate degradation | 3 / 4 | 0.030 |
|  | Glyoxylate and Dicarboxylate Metabolism | 4 / 7 | 0.040 |
|  | Nitrogen metabolism | 4 / 7 | 0.040 |
|  | Fructose and mannose metabolism | 5 / 10 | 0.041 |
|  | Arginine and Proline Metabolism | 13 / 38 | 0.048 |
| HILIC | Glycerophospholipid metabolism | 16 / 42 | <0.001 |
|  | Glycosphingolipid metabolism | 8 / 22 | 0.007 |
|  | Carnitine shuttle | 9 / 28 | 0.011 |
|  | Fatty acid activation | 9 / 28 | 0.011 |
|  | De novo fatty acid biosynthesis | 11 / 38 | 0.013 |
|  | Xenobiotics metabolism | 6 / 18 | 0.024 |
|  | Vitamin B9 (folate) metabolism | 3 / 6 | 0.026 |
|  | C21-steroid hormone biosynthesis and metabolism | 9 / 36 | 0.046 |
|  | Fatty Acid Metabolism | 7 / 26 | 0.046 |
| Metabolic pathways significantly associated with time above range (glucose level > 140 mg/dL) during the first 48 hours of surgery. Pathway enrichment analysis was performed using Mummichog for C18 and HILIC chromatography. Pathway size refers to the total number of putative compounds associated with a given pathway, while overlap size represents the number of significant putative compounds detected. P-values indicate statistical significance of pathway enrichment. | | | |

**Supplementary Table 7.** Pathways Associated with Surgical Stress in the Whole Cohort

| **Column** | **Pathway** | **Overlap size /**  **Pathway size** | **p-value** |
| --- | --- | --- | --- |
| C18 | Arachidonic acid metabolism | 26 / 29 | 0.002 |
|  | C21-steroid hormone biosynthesis and metabolism | 41 / 50 | 0.006 |
|  | Prostaglandin formation from arachidonate | 36 / 44 | 0.010 |
|  | Glycerophospholipid metabolism | 35 / 43 | 0.012 |
|  | Squalene and cholesterol biosynthesis | 8 / 8 | 0.018 |
|  | Androgen and estrogen biosynthesis and metabolism | 21 / 25 | 0.023 |
|  | Porphyrin metabolism | 14 / 16 | 0.029 |
|  | Leukotriene metabolism | 22 / 27 | 0.037 |
|  | Bile acid biosynthesis | 30 / 38 | 0.038 |
|  | Methionine and cysteine metabolism | 19 / 23 | 0.039 |
|  | Purine metabolism | 42 / 55 | 0.045 |
| HILIC | Carnitine shuttle | 22 / 28 | 0.021 |
|  | Urea cycle/amino group metabolism | 36 / 49 | 0.032 |
|  | Glutathione Metabolism | 5 / 5 | 0.036 |
|  | Bile acid biosynthesis | 22 / 29 | 0.041 |
| Metabolic pathways showing significant temporal variation across four time points in the whole cohort. Pathway enrichment analysis was performed using Mummichog for C18 and HILIC chromatography. Pathway size refers to the total number of putative compounds associated with a given pathway, while overlap size represents the number of significant putative compounds detected. P-values indicate statistical significance of pathway enrichment. | | | |

**Supplementary Table 8.** Pathways Associated with Stress Hyperglycemia.

| **Column** | **Pathway** | **Overlap size /**  **Pathway size** | **p-value** |
| --- | --- | --- | --- |
| C18 | Squalene and cholesterol biosynthesis | 6 / 8 | 0.007 |
|  | Bile acid biosynthesis | 17 / 38 | 0.013 |
|  | Vitamin B9 (folate) metabolism | 4 / 5 | 0.018 |
|  | Glycerophospholipid metabolism | 18 / 43 | 0.021 |
|  | Putative anti-Inflammatory metabolites formation from EPA | 5 / 8 | 0.034 |
|  | Propanoate metabolism | 4 / 6 | 0.043 |
|  | Histidine metabolism | 11 / 25 | 0.043 |
| HILIC | Glycosphingolipid metabolism | 12 / 22 | <0.001 |
|  | Glycerophospholipid metabolism | 14 / 42 | 0.006 |
|  | Leukotriene metabolism | 9 / 23 | 0.007 |
|  | C21-steroid hormone biosynthesis and metabolism | 12 / 36 | 0.010 |
|  | Arachidonic acid metabolism | 6 / 17 | 0.036 |
|  | Vitamin B9 (folate) metabolism | 3 / 6 | 0.038 |
|  | TCA cycle | 3 / 6 | 0.038 |
| Metabolic pathways showing significant stress hyperglycemia group difference in temporal variation across four time points. Pathway enrichment analysis was performed using Mummichog for C18 and HILIC chromatography. Pathway size refers to the total number of putative compounds associated with a given pathway, while overlap size represents the number of significant putative compounds detected. P-values indicate statistical significance of pathway enrichment. | | | |

**Supplementary Table 9**. Top 20 Metabolites Ranked by Feature Importance

| **Rank** | **Column** | **m/z_RT (s)** | **Adducts** | **Mummichog, KEGG Label** | **Abbreviation** | **Relevance** |
| --- | --- | --- | --- | --- | --- | --- |
| 1 | C18 | 381.204_171.1 | *[M+^37^Cl]^−^* | 4-hydroxy-all-trans-retinyl acetate | 4-OH-Retinyl Acetate | Proteomic analyses of human and guinea pig heart failure were consistent with a decline in levels of cardiac all-trans retinoic acid (ATRA).^12^ In addition, the presence of ATRA improved insulin sensitivity and decreased the retinol to RBP4 ratio (p < 0.05).^13^ |
| 2 | HILIC | 170.0842_90.6 | *[M−H_2_O−H]^−^* | *1-Pentanesulfonic acid | PA | This compound is a part of the exposome and is not endogenous to humans. |
| 3 | HILIC | 281.0898_74.1 | *[M−CH_3_COO]^−^* | Salsolinol 1-carboxylate | Sal-1-COOH | Salsolinol has shown to inhibit inflammation in angiotensin II induced Human Cardiac Fibroblasts (HCFs).^14^ Similarly, Salsalate lowers HbA1c levels while also improving other markers of glycemic control for T2D.^15^ |
| 4 | HILIC | 317.2128_25.4 | *[M−H_2_O−H]^−^* | 5-Hydroperoxy-6-trans-8,11,14-cis-eicosatetraenoate | 5-HpETE | - |
| 5 | HILIC | 225.0294_17.8 | *[M+ACN−H]^−^* | O-Phospho-L-serine | P-Serine | One paper found that in a population of hospitalized patients at the Peking University First Hospital, those who had higher serine concentrations had lower risk of coronary heart disease.^16^ In addition, L‐serine concentration has been found to be significantly altered in diabetes, including one study in children with T1D which found that the concentration of L‐serine in plasma was decreased by 42% in the T1D patients.^17^ |
| 6 | C18 | 315.198_142.7 | *[M−H_2_O−H]^−^* | Prostaglandin A2 | PGA2 | Prostaglandin I, a related molecule, was produced endogenously during myocardial ischemia-reperfusion injury. It has also been shown to have a protective effect on cardiomyocytes.^18^ In addition, PGA2 was found to have an insulin sensitizing effect when in the presence of nuclear receptor 4A3 (NR4A3).^19^ |
| 7 | C18 | 434.1588_138.9 | *[M−H_2_O−H]^−^* | Alpha-CEHC-glucuronide | α-CEHC-gluc | Levels of alpha-CEHC-glucuronide were found to be elevated in urine samples of patients with T1D, which could hint at metabolic oxidative stress.^20^ No information on cardiac outcomes found. |
| 8 | HILIC | 508.376_35.6 | *[M−H]⁻* | 1-Alkyl-2-lyso-sn-glycero-3-phosphocholine | Lyso-PAF | No information relevant to the outcomes of interest were found. |
| 9 | C18 | 195.139_144.7 | *[M−2H]^2−^* | Deoxycholic acid | DCA | See Table 7 |
| 10 | C18 | 159.1026_297.2 | *[M−H+O]^−^* | Octanoic acid | C8:0 | Perfluoro–octanoic acid (PFOA) targets platelet membranes, whose accumulation was associated with increased membrane fluidity, increased activation at resting condition, and calcium uptake/activation aggression, all of which led to increased cardiovascular risks.^21^ In addition, octanoic acid found to have inverse relationships with diabetes risk. This was enhanced for people with higher genetic risk factors for diabetes.^22^ |
| 11 | C18 | 91.0287_20.7 | *[M−2H]^2−^* | 3-Methoxy-4-hydroxyphenylethyleneglycol | MHPG | No information relevant to the outcomes of interest were found. |
| 12 | HILIC | 730.3354_36.6 | *[M−H_2_O−H]^−^* | Beta-casomorphin (1-6) | β-CM(1–6) | No information relevant to the outcomes of interest were found. Most research is on b-casomorphin 7. |
| 13 | C18 | 204.0858_23.8 | *[M+ACN−H]^−^* | 6-Deoxy-L-galactose | 6-Deoxy-Gal | D‐galactose induces an aging heart, which results in increased oxidative stress and reduced levels of antioxidants.^23^ No information on glycemic outcomes was found. |
| 14 | C18 | 677.2446_148.7 | *[M+K−2H]^−^* | 10,11-dihydro-12R-hydroxy-leukotriene C4 | 12R-OH-LTC4 | See Table 6. |
| 15 | C18 | 351.0932_134.7 | *[M−H_2_O−H]^−^* | 4-(2-amino-3-hydroxyphenyl)-4-oxobutanoic acid O-glucoside | 4-AHPBA-Glc | No information relevant to the outcomes of interest were found. |
| 16 | HILIC | 169.0089_271.8 | *[M−H]⁻* | Diethylthiophosphoric acid | DETP | DEHP exposure has been shown to modify cardiac electrical conduction in rats, including a decrease in heart rate and prolongation of PR/QT intervals.^24^ In addition, increased levels of DETP in urine samples were found to be associated with observed insulin resistance.^25^ |
| 17 | HILIC | 322.042_74.9 | *[M−H]⁻* | CMP | - | See Table 7 |
| 18 | C18 | 155.0462_90.0 | *[M−H+O]^−^* | 3-Methylimidazoleacetic acid | 3-MIAA | 3-Methylimidazoleacetic acid is a breakdown product of histamine.^26^ Pancreatic function is intricately linked to histamine levels – greater insulin levels have been associated with decreased histamine in rats with diabetes.^27^ In addition, Histamine may contribute to atherosclerosis development by increasing the amount of H1R mRNA.^27^ |
| 19 | HILIC | 124.087_121.0 | *[M−H]⁻* | N-Methylhistamine | NMH | No information relevant to the outcomes of interest were found. |
| 20 | C18 | 311.2955_299.1 | *[M−H]⁻* | Icosanoic acid; Eicosanoic acid; Arachidic acid$Phytanate; Phytanic acid | C20:0, Phy | Very general, says that "extensive lines of evidence imply a diverse and disease-specific contribution of individual AA metabolites to cardiovascular health and complications".^28^  AA can prevent whole-body insulin resistance that is induced by high-fat diets.^29^ |
| Top 20 metabolites identified by feature importance analysis from the ridge logistic regression model for predicting stress hyperglycemia. Metabolites are ranked by their contribution to model performance, with chromatographic details (column type, m/z, retention time), chemical annotations from Mummichog and KEGG, and abbreviations provided. The “Notes” column indicates metabolites that overlap with significant findings from previous analyses: one metabolite (rank 14) was also identified in the whole cohort analysis (Table 6), and three metabolites (ranks 9, 17, 18) were identified in the group difference analysis (Table 7). *Metabolite identified by manual annotation. | | | | | | |
